# *Fusobacterium nucleatum* present in the saliva of oral cancer subjects can activate niche defense of oral squamous cell carcinoma

**DOI:** 10.1101/2023.10.10.561552

**Authors:** Partha Jyoti Saikia, Lekhika Pathak, Shirsajit Mitra, Tutumoni Baishya, Rupam Das, Ibrahim S Akeel, Bidisha Pal, Bikul Das

**Author notes:** Corresponding Author: Bikul Das, MD, PhD.

## Abstract

Oral cancer is a subset of head and neck cancer (HNC), has a high incidence rate in this malignancy group. Cancer Stem Cells (CSCs) are population of the heterogeneous malignant cells present within oral tumor microenvironment. CSCs’ stemness permits them to control several signaling pathways and so play a role in cancer progression and relapse. A number of studies have recently demonstrated the presence of specific oral bacteria populations and their lipopolysaccharides (LPS) in the tumor microenvironment. The precise mechanism of action in the initiation, progression, and relapse of oral cancer by the oral bacteria are yet to be determined. We previously reported pathogenic bacterial internalization in CSCs. Based on the findings; we have developed an in-vitro model to investigate how oral microbiota may integrate into the tumor microenvironment’s CSC population and control its activity. Notably, we found that live bacteria and their LPS, mostly *Fusobacterium nucleatum* isolated from clinical subjects, were capable of invading CSCs in the *in-vitro* experimental design setup. Post the host-pathogen interaction; it enabled the activation of a niche modulatory tumor stemness defense (TSD) phenotype in the CSCs. These aggressive CSCs with the TSD phenotype have been found to have a critical role in the progression and relapse of oral cancer.

## Introduction

Oral squamous cell carcinoma (OSCC) is one of the most common form of Head and neck cancer and a very aggressive cancer (1). One of the key reasons of aggressiveness is the presence of cancer stem cells (CSCs) in OSCC. Cancer stem cells are subpopulation of cells present in tumor and are responsible for tumor initiation, recurrence, metastasis, and therapeutic resistance. A defining characteristic of CSCs is “stemness,” which is responsible for their ability to self-renew as well as its role in tumor progression. Stemness in CSCs are maintained by integrated gene network system (2). Inflammatory mediators regulate cancer stemness genes in several cancer types. Notably, in oral squamous cell carcinoma (OSCC), both HIF-1α and NF-kB play pivotal roles in sustaining cancer stemness(3). Furthermore we have shown that up regulation of stemness genes in oral CSCs are maintained by MYC dependent HIF 2 α stemness pathway (4).

CSCs reside in specific niches in the tumor microenvironment. During chemotherapy or due to oxidative stress or due to pathogen invasion, CSCs exhibit a unique defense mechanism. Importantly the defense mechanism reprograms the CSCs to a highly tumorigenic phenotype i.e the Tumor stemness defense phenotype (TSD) (5). Recently we reported that preferential invasion of pathogens such as *Mycobacterium bovis* and Mtb18b in post hypoxia side population cells Spm(hox) +, ABCG2+ CSCs of seven cell lines including SCC 25 (representative cell line of OSCC) exhibited a niche defense mechanism. The TSD phenotype defended their niches by expression of p53/HMGB1 complex. Importantly this complex can be used to target CSCs residing in their niches by pathogen induced bystander apoptosis (PIBA) (5). However pathogen induced niche defense mechanism with high tumorigenic potential can oblige the cancer to be aggressive. Therefore there is a need to study the CSC niche defense and how TME can activate this niche defense.

Recent studies reported significant expression of genes involved in bacterial LPS biosynthesis in tumor sites of OSCC patients (6). Experimental models have demonstrated bacterial endotoxin LPS involved in carcinogenesis (7). In an *in-vitro* model, it has been reported that the LPS from the oral bacterium *Fusobacterium nucleatum* activates inflammatory cytokines like IL-6 and TNF-α, potentially influencing cancer development (8). Additionally, another study reported that the LPS from the oral bacterium *Porphyromonas gingivalis* triggers a Toll-like receptor 4 (TLR4) response, which inhibits apoptosis and promotes tumor cell proliferation and invasiveness(8). Notably in OSCC, LPS can mediate cancer stemness by epithelial mesenchymal transition via TLR 4 expression (9). Microbe induced TLR expression in oral cancer cells may increase NF-kB which in turn promotes cancer stemness (10). Thus, the studies have confirmed the role LPS in bacteria mediated cancer stemness in oral cancer. In this context there is a strong need to develop clinical and in vitro models to study LPS of oral microbiota induced CSC niche defense.

Saliva microbiome profiles of OSCC patients have shown high abundance of pathogenic bacteria compared to healthy controls (11). Pathogens such as FN bind to the epithelial cells by Fusobacterium adhesin A (FadA) and lipopolysaccharides LPS (bacterial endotoxin). Fad A and LPS are recognized by TLRs (12). These TLRs in activates chronic inflammatory pathways which in turn incudes stemness.

The current standard treatment approach for managing locally advanced, unresectable head and neck cancer, which includes oral cancer, involves a combination of chemoradiation with cisplatin. However, nearly 60% subjects do not respond to platinum therapy. In our findings, it was observed that cisplatin treatment has the potential to trigger the TSD phenotype in osteosarcoma and several other cancer types. This effect is achieved through the activation of the VEGF-HIF1α pathway (13). Furthermore, in the context of oral cancer, cisplatin treatment was found to induce the expression of Bmi-1 and activate the IL-6/STAT3 pathway. Notably in the clinical setting, we also found the presence *Fusobacterium nucleatum* in primary tumor derived EpCAM+/ABCG2+ CSCs of platinum treated patients (14). Thus there is a possibility that platinum treated patients having TSD phenotype harboring FN might show resistance to the therapy. Therefore there is need to develop pre-clincial model to screen saliva of OSCC patients with ability to induce TSD phenotype, so that we can identify patients who may or may not respond to cisplatin therapy.

Hence, here we have developed an *in-vitro* assay of culturing SCC-25 cells with saliva derived LPS, or direct co-culture of salivary cell aggregates and SCC-25 cells. In this manner, we have identified that FN contain in Saliva of aggressive OSCC patients can modulate the TSD phenotype.

## Materials and Method

### SCC-25 cell line culture

All the necessary experimental procedures were undertaken inside BSC-class II facility in accordance with guidelines of “Institutional Bio-safety Committee” of KaviKrishna Laboratory. The SCC-25 cell line obtained from ATCC (CRL-1628) was cultured in a 1:1 mixture of Dulbecco’s modified Eagle’s medium and Ham’s F12 medium containing 1.2 g/L sodium bicarbonate, 2.5 Mm L-glutamine, 15 Mm HEPES and 0.5 Mm sodium pyruvate and supplemented with 400 ng/ml hydrocortisone and 10% fetal bovine serum (FBS). The SCC-25 cells were counted via the trypan blue exclusion method and the cells (∼1 × 105) were seeded in 6-well tissue culture plates and incubated overnight at 37℃ in the 5% CO2 incubator and then the cells were subjected to various assays.

### Bacterial lipopolysaccharide (LPS) treatment

Following two rounds of rinsing with phosphate-buffered saline (PBS) from Himedia Mumbai, India, the cells in each well were subjected to treatment with 1,000 ng/mL of LPS obtained from Sigma, USA, following a previously described protocol (15). Subsequently, after a 3-day incubation period, the cells were once again washed with PBS, and a fresh DMEM medium was introduced. The cells were then allowed to grow for a duration of one week.

### Patient selection

The study was approved by the ethical committee at KaviKrishna Laboratory, IIT Guwahati Research Park, Indian Institute of Technology, Guwahati. Oral cancer patients (n=22, table 1) and four healthy individuals were enrolled in the study. Following written informed consent, saliva was collected from these individuals. The clinical cases with head & neck cancer were selected based on history, and diagnosis performed at Guwahati Medical College. Previously treated subjects of oral squamous cell cancer were recruited based on pathological data.

### Saliva collection and processing

Saliva collection was performed using the spitting method, as previously outlined (16). In the early morning, patients gathered saliva from the floor of their mouths and expelled it into a collection tube at 60-second intervals. Approximately 1 ml of saliva was collected, dissolved in DMEM media within a 15 ml tube, and transported on ice to the laboratory. Upon arrival, the saliva sample was promptly centrifuged at 10,000 rpm for 10 minutes at 4°C. The resulting pellet was then resuspended in 1 ml of DMEM, and each flask was treated with 100 μl of the processed saliva. Following a 3-day incubation period, each flask underwent treatment with amikacin and metronidazole for 24 hours to eliminate any remaining extracellular bacteria. Subsequently, the cells were cultured for a week.

### Ciprofloxacin treatment of the saliva treated and LPS treated SCC-25 cell line

10 μg/ml Ciprofloxacin was added to the processed saliva and LPS for 24 hours to make them sterile. Then after adding this saliva and LPS to the SCC-25 cells, we continued to add 1μg/ml of ciprofloxacin and metronidazole for 7 days. This was done because the cell lines contained *F. nucleatum* and they were sensitive to metronidazole and ciprofloxacin. Ciprofloxacin has been shown to be effective for intracellular and extracellular bacteria (21). Among the patients 18-22, the 21 and 22 contained *F. nucleatum* as well as *Staphylococcus*, and *Streptococcus* infection, and these bacteria were sensitive to ciprofloxacin as well as per the bacterial culture and sensitivity reports.

### Immunomagnetic sorting of ABCG2+ cells

The ABCG2 cells were isolated using an immunomagnetic sorting method that we recently employed to purify CD271+ cells. This method involved the use of phycoerythrin (PE) conjugation, as described previously (17). To facilitate this process, the ABCG2 antibody (#ab 3380, Abcam) was initially conjugated with PE using the SiteClick antibody labelling kit, as previously described (17). We achieved a purity level of 94% through this approach. Additionally, we utilized a biotin-conjugation method, where the ABCG2 antibody was conjugated with biotin (#ab95692, Abcam) and subjected to biotin-positive selection (#18559, Stem Cell Technologies, BC). This method yielded a similar level of enrichment as the PE conjugation method.

### Real-time PCR

Total RNA was extracted from the cell lines one week after treatment with either LPS or saliva, using Trizol reagent (ThermoFisher Scientific, # 15596018). Subsequently, cDNA was synthesized through reverse transcription utilizing M-MLV reverse transcriptase from BIO RAD. Quantitative polymerase chain reaction (qPCR) was conducted in a Qiagen Rotor gene Q, following the previously described protocol (18). The qPCR process involved utilizing 2 ng of initial cDNA and running 50 cycles at 94°C for 60 seconds. To ensure accurate measurements, RNA levels were normalized to the reference gene glyceraldehyde 3-phosphate dehydrogenase (GAPDH) and quantified using the ΔΔCt method with SDS software, version 2.2.1 (Applied Biosystems, Foster City, USA). TaqMan gene expression primers for the following genes were employed: HIF-2α (Hs01026149_m1), Nanog (Hs02387400_g1), Oct 4 (Hs03005111_g1), ABCG2 (Hs00184979_m1), Sox2 (Hs00602736_s1), MYC (Hs00153408_m1), p21 (Hs00355782_m1), and GAPDH (Hs00266705_g1).

### Clonogenic assay

The assay was carried out following the procedures as described previously (18, 19). In brief, a total of 1×10^3 freshly sorted cells, including ABCG2+ and ABCG2-populations, were placed in methylcellulose medium. This medium was prepared using either Methocult M3134 from Stem Cell Technologies, BC, or Methylcellulose powder (# AC182312500, Fisher Scientific) that was reconstituted in RPMI media with 10% FBS at KaviKrishna Lab, Guwahati, India. These cells were seeded into a 6-well plate, then incubated at 37°C with 5% CO2, and the colonies were counted after a two-week incubation period.

### Western blot

Western blot was performed as previously described (18). Briefly, protein extracts were obtained using the cell lysis buffer (Sigma Aldrich) mixed with protease inhibitor cocktail (Sigma Aldrich). 50 μg of the clear protein lysate was resolved on 4% to 12% SDS-PAGE gels and transferred to 0.45-μm nitrocellulose membranes (Bio-Rad, CA, USA). The blocked membranes were incubated overnight with the following antibodies: HIF-2α (Novus Biologicals, Littleton, CO), MYC, Nanog, Oct-4, SOX-2 and β-actin (Cell Signaling technology). Immunoblotting was detected by enhanced chemiluminescence (Thermo Scientific, Rockford, IL, USA) according to the manufacturer’s instructions. Densitometry study was done using the Α View software (Α InfoTech Α Imager 3400 Light Cabinet Camera Transilluminator Imaging UV, CA, USA). Following antibodies were used: Hm-MYC, Hm-HIF-2α, Hm-Nanog, Hm-Oct-4, and Hm-SOX-2 (Cell Signaling technology). Immunoblotting was detected by enhanced chemiluminescence (Thermo Scientific, Rockford, IL, USA) according to the manufacturer’s instructions. Densitometry study was done using the ΑView software (Α Innotech Α Imager 3400 Light Cabinet Camera Transilluminator Imaging UV, CA, USA).

### ELISA

To determine the protein levels of HIF-2α, MYC, Nanog, Oct-4, and SOX-2, we employed In Cell ELISA with a horseradish peroxidase (HRP)-conjugated detection reagent, specifically using the In-Cell ELISA Colorimetric detection kit (Thermo Fisher, #62200). Additionally, we utilized standard ELISA kits as previously described (19). The HIF-2α antibody, designated as MBS-702348, was acquired from MyBiosource, located in San Diego, CA.

### Matrigel invasion assay

A Boyden chamber invasion assay was conducted in accordance with previously published procedures (19). To outline the process briefly, we used polyvinyl membrane-based chambers with 8-μm pore sizes (Corning Life Sciences, Lowell, MA), which were pre-coated with 100 μl of ice-cold Matrigel (7.5 mg/ml; BD Biosciences, San Diego). These coated chambers were then incubated at 37°C for 4 hours. Following trypsin neutralization, the appropriate number of cells was added to the upper chamber, while the lower chamber was filled with the required media. The chamber was then incubated at 37°C for a duration of 8 to 24 hours, and the invading cells were subsequently counted after staining with crystal violet (20). If needed, the invading and non-invading cells were isolated through brief trypsinization and expanded using the appropriate culture medium.

### PCR

The PCR was run for 32 cycles on a Peltier thermal cycler (PTC-200 DNA engineTM, MJ Research Inc., Watertown, MA, U.S.A.) as described previously (22). Briefly, each cycle consisting of denaturation at 94°C for 1 min, primer annealing at 68oC for 30 sec, and extension at 72°C for 1 min. The final cycle has an additional 10 min extension at 72°C. A 2 ml aliquot of the reaction mixture was analyzed by 1.5% agarose gel electrophoresis in a Tris-acetate buffer (0.04 M Tris-acetate, 0.001 M EDTA, and (pH 8.0) at 100 V for 30 min. The amplification products obtained by the *F. nucleatum* were stained with ethidium bromide and visualized by UV transillumination. The following primers were used: conserved forward primer, 5’-CGG GAG GCA GCA GTG GGG AAT-3’; and *F.nucleatum* reverse primer, 5’-TTG CTT GGG CGC TGA GGT TC.

### *F. nucleatum* culture and infection of SCC-25 cell line

The *F. nucleatum* obtained from ATCC (#25586) was cultured anaerobically for 24 hours in brain heart infusion broth supplemented with yeast extract (5 mg/ml), hemin (5 μg/ml) and menadione (1 μg/ml) as described previously (23). The culture is then washed with PBS and the SCC-25 cell line was infected with *F. nucleatum* at a multiplicity of infection of 1:50, quantified using optical density (OD600), and incubated at 37°C with 5% CO2 incubator for 12 hours.

### Immunomagnetic sorting of ABCG2+ cells

The *F. nucleatum* treated SCC-25 cell line culture was subjected to immunomagnetic sorting for ABCG2+ cells as described previously.

### Real time PCR

Total RNA was extracted, and qPCR was performed as described previously. The following TaqMan gene expression primers used were HIF -2α (Hs01026149_m1), Nanog (Hs02387400_g1), Oct 4 (Hs03005111_g1), ABCG2 (Hs00184979_m1), Sox2 (Hs00602736_s1), MYC (Hs00153408_m1), p21 (Hs00355782_m1), and GAPDH (Hs00266705_g1). The following primers were used to detect specificity for 16S Rrna of *F. nucleatum*: 5′-CCGCGGTAATACGTATGTCACG-3′ and 5′-TCCGCTTACCTCTCCAGTACTC-3′.

### Western blot

Western blot was performed as previously described. The following antibodies were used: Hm-MYC, Hm-HIF-2α, Hm-Nanog, Hm-Oct-4, and Hm-SOX-2 (Cell Signaling technology). Immunoblotting was detected by enhanced chemiluminescence (Thermo Scientific, Rockford, IL, USA) according to the manufacturer’s instructions. Densitometry study was done using the ΑView software (Α Innotech Α Imager 3400 Light Cabinet Camera Transilluminator Imaging UV, CA, USA).

### Statistical analysis

The statistical calculations were performed with GraphPad Prism 4.0 (Hearne Scientific Software) using Student t test and One-Way ANOVA with Dunnett post-hoc test.

## Results

### Bacterial load in OSCC patient derived saliva

To study the oral microbiota induced stemness, we first obtained saliva of n=25 OSCC patients of grade II, III and IV or loco-regional recurrent OSCC. The Saliva was processed to obtain DNA and also LPS. The DNA was extracted from 1 ml of saliva to estimate the 16sRNA by qPCR, whereas, LPS was obtained from ml of Saliva by using a LPS extraction kit (Sigma). We found that among the 25 subjects, the bacterial load was highest in six subjects (∼8×10^8^/ml vs 1×10^9^/ml), with the corresponding high concentration of LPS (∼2-6-ug/ml). Notably, LPS was extracted from the saliva (10 ml) of n=25 OSCC patients. Only 2 patients (#4 and 5 patients) showed the presence of >1mg of LPS in 10 ml. Thus, we did not find any significant association between the tumor staging/relapse and the bacterial load or LPS content.

### LPS of OSCC Patient derived saliva induces inflammatory cytokines of SCC-25 cell

We proposed that LPS obtained from patients with high-grade and relapsed OSCC may exhibit higher potency in inducing inflammation of blood cells than LPS of low-grade OSCC patients. To test this hypothesis, the saliva derived LPS (1ug/ml) was then used to treat donor-derived PBMC for 72 hours followed by the measurement of IL-8. In this manner, we found that high grade and relapsed OSCC patients derived LPS (patients 6, 13, 16, 20-23) showed the highest increase of IL-8 expression (4-6 fold increase; Figure 1) as compared to untreated PBMC. These results indicate that LPS obtained from high-grade tumors exhibited potent inflammatory activity.

**Figure 1.**
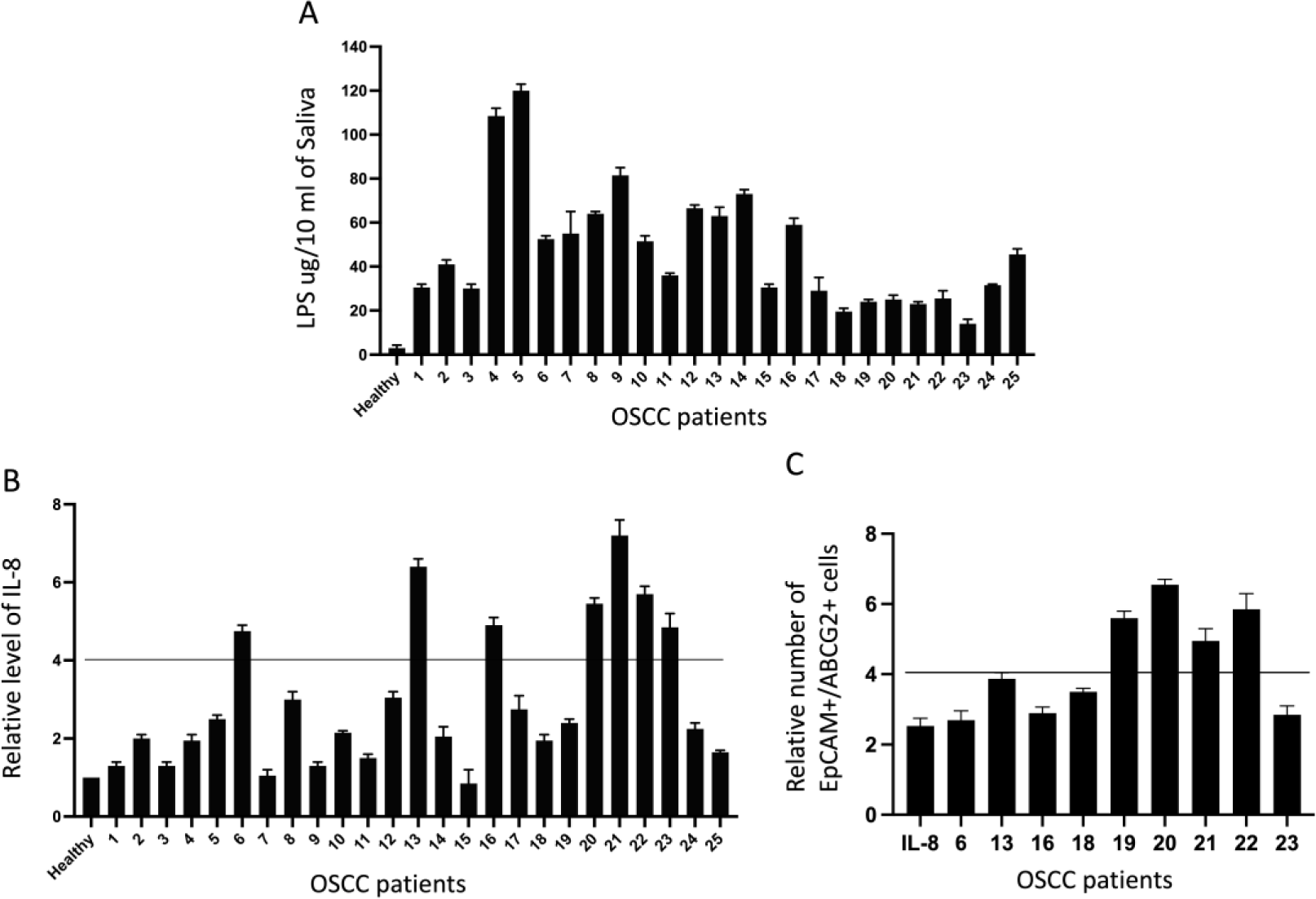
Inflammatory cytokines present in patient-derived LPS. **A**. LPS levels isolated from saliva samples of patients. **B.** Measurement of IL-8 in the peripheral blood mononuclear cells (PBMC) of healthy donors treated with patient’s saliva-derived LPS. **C**. The EpCAM+/ABCG2+ CSCs found in the patient tumor sample. Data is the mean of three independent experiments. Dash line represents the clinically significant increase in the levels of IL-8 and EpCAM+/ABCG2+ cells respectively.

### Salivary lipopolysaccharide (LPS) induces tumor stemness defense(TSD) phenotype in SCC-25 derived ABCG2+ CSCs

We proposed the TSD phenotype as a stress-response of cancer stem cell (CSCs) to defend their niches. Therefore, we reasoned that the saliva of OSCC patients with poor prognosis may be enriched in LPS capable of inducing TSD phenotype in OSCC cell lines. To investigate this possibility, first we used LPS of patients 6, 13, 16, 18-23 to treat the SCC-25 cell line. We have recently found that the SCC-25 cell line maintain a dormant population of EpCAM+/ABCG2+ CSCs which reprogram to TSD phenotype to defend the CSC niche against hypoxia/oxidative stress or platinum induced stress. Thus, the SCC-25 cell line was treated with LPS (1ug /ml) for 3-days (Figure 2A) and then cells were cultured for a week in the serum-free medium, and then immunomagnetic sorting was done to evaluate the expansion of EpCAM+/ABCG2+ CSCs. As a control, cells were treated with 20 ng/ml IL-8 for 3 days. After a week, we found that LPS of patients 19-22 showed the 4-5-fold increase of EpCAM+/ABCG2+ cells (Figure 2A), whereas IL-8 showed only 2-fold increase of EpCAM+/ABCG2+ cells (data not shown). Hence, we used the LPS of patients 19-22 to evaluate for the TSD phenotype (Figure 2 A-G). We recently characterized the TSD phenotype in OSCC derived EpCAM+/ABCG2+ cells with higher stemness gene expression, transient expansion of ABCG2+CSCs, higher clonogenicity, invasive ability and niche modulatory activity. Indeed, the stemness genes HIF-2α, Nanog, Oct4 and Sox2 were significantly increased by 2-3-fold (p<0.01) in LPS treated SCC-25 cells versus untreated SCC-25 cells (Figure 2B), and the associated increase of EpCAM+/ABCG2+ CSCs, and their TSD phenotype. Thus, the proliferation and clonogenic assay revealed a 4-fold increase in both proliferation and clonogenic activity of LPS treated versus untreated ABCG2+CSCs CSCs (Figure 2D-E). We noted that LPS treatment had no effect on clonogenic activity and expansion in the EpCAM+/ABCG2-CSC population (Figure 2F). Importantly, real-time PCR assay showed 2-4 fold increase of TSD phenotype associated genes in LPS treated vs untreated EpCAM+/ABCG2+CSCs (Figure 2G).

**Figure 2.**
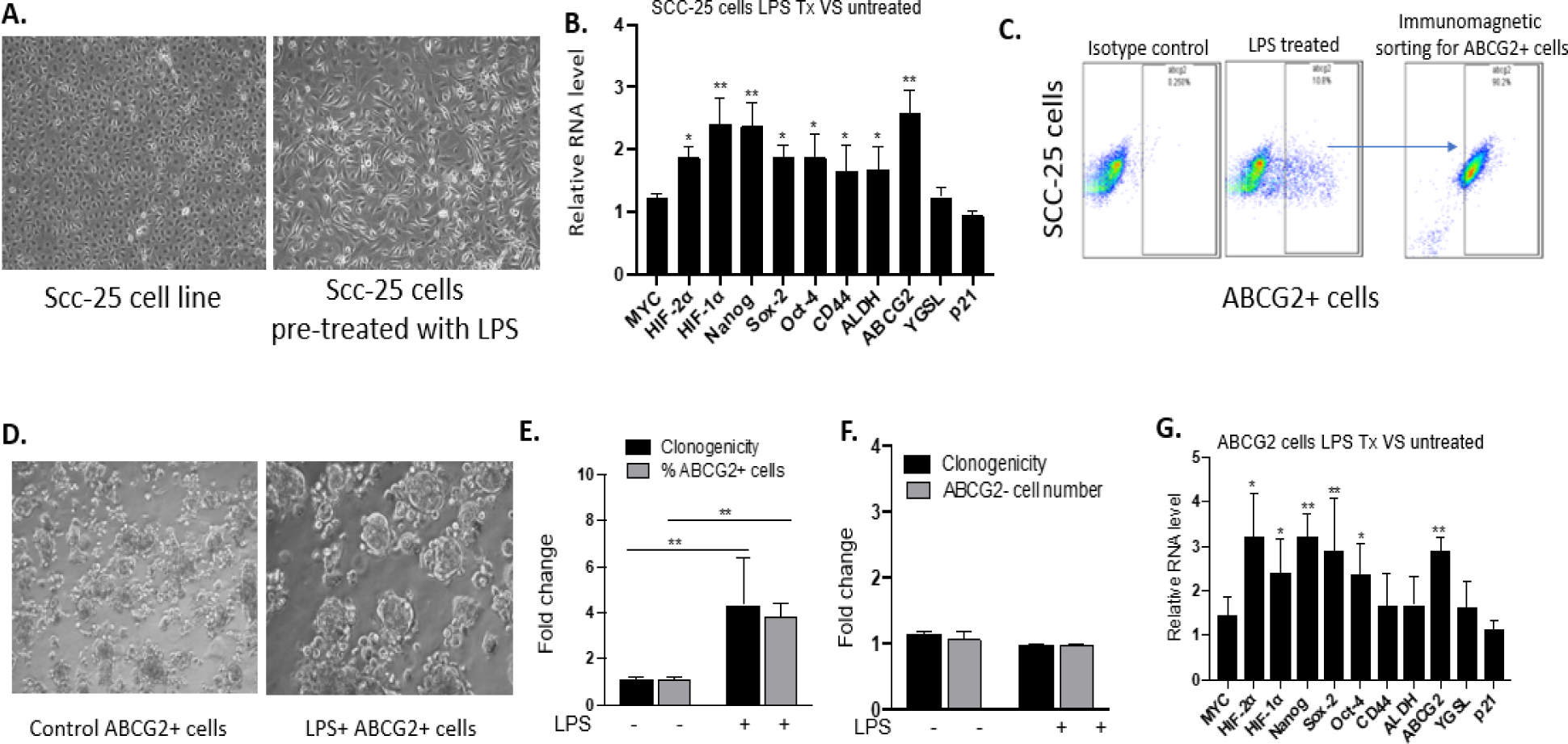
LPS induced tumor stemness switch in SCC-25 cell line. A) Representative phase contrast image of SCC-25 cells with and without LPS (patient #22 derived) treatment. B) qPCR analysis shows upregulation of stemness-associated genes involved in the HIF-2α stemness pathway in SCC-25 cells post-LPS treatment (patients #19-22). C) Representative flow cytometry shows further enrichment of ABCG2+ cells post LPS treatment (#patient 20). D-F) LPS treatment enhanced Clonogenic potential and ABCG2+ cell number. Additionally, the invasive and niche modulatory capacity of ABCG2+ vs ABCG2-cells was enhanced. G) qPCR analysis of immunomagnetically sorted ABCG2+ cells shows increased expression of genes involved in the HIF-2α stemness pathway. ABCG2+ cells were obtained from the immunomagnetic sorting population of EPCAM+ cells of SCC-25. N = 3 independent experiments; error bar represents mean ± SEM. *p<0.05, **p<0.001, ANOVA.

Next, we performed Matrigel invasion assay as previously described (19) to evaluate the invasive potential of EpCAM+/ABCG2+CSCs in the LPS-treated versus untreated group. We found significantly higher number of invading ABCG2+ CSCs (invading cells/10^4^ total cells) in the treated versus untreated group (Figure 3A-B). These results suggest that LPS obtained from relapsed OSCC subjects may induce TSD phenotype in SCC-25 cells.

**Figure 3.**
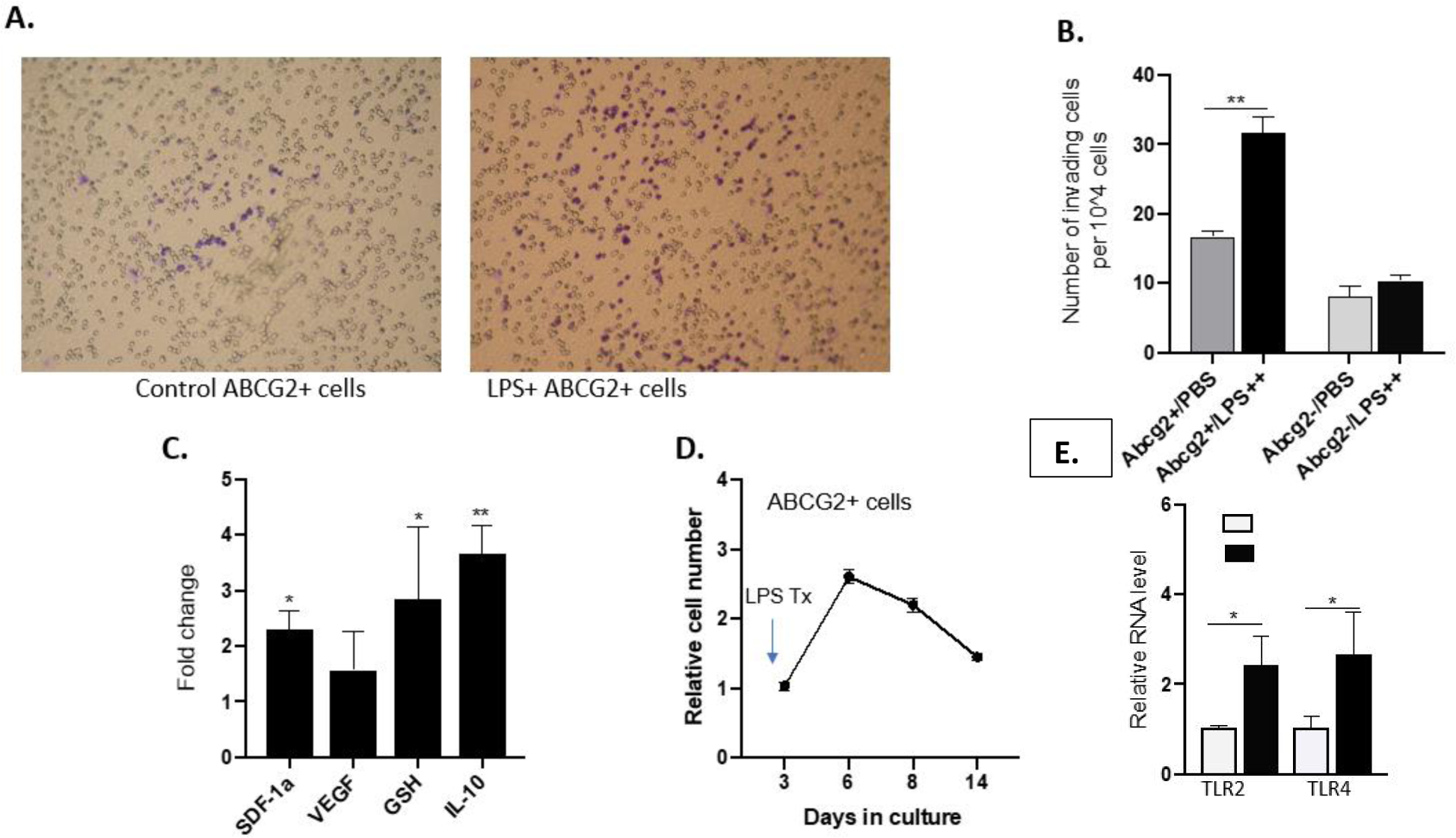
LPS treated ABCG2+ cells exhibit tumor stemness switch. A. Representative microscopic images of the Matrigel invasion assay show increased number of cells passing through the pores post LPS treatment. B) Comparison between the relative number of invading cells in LPS treatment group and PBS treatment group. Note the increased number is limited to the ABCG2+ cell population. C) Condition media of ABCG2+ CSCs exhibit niche modulatory activity. D) ABCG2+ cell number increased transiently and lasted for approximately two weeks. E) day 7 after LPS treatment. qPCR analysis indicates that the LPS treatment led to increased expression of TLR2/4 genes. N = 3 independent experiments; error bar represent mean ± SEM. *p<0.05, **p<0.001, ANOVA.

To further assess the niche modulatory potential, we obtained the conditioned media (CM) from the ABCG2+ CSCs and treated with the mesenchymal stem cells (MSCs) for 2 weeks. MSCs are integral part of the tumor microenvironment (TME) and are known to secrete growth factors important for tumor growth, such as stromal derived factor 1 α (SDF-1 α), vascular endothelial growth factor (VEGF), glutathione, and interleukin 2 (IL-2) (24, 25). Notably, we found that the MSCs treated with the conditioned media of ABCG2+ CSCs exhibited 2 fold increased secretions of SDF-1 α,1 fold increased in VEGF, 3 fold increased in glutathione (GSH 4), and 4 fold increased in IL-10 (Figure 3C). These results suggest that LPS treated EpCAM+/ABCG2+ CSCs exhibited nihce modulatory activity.

Recently, using a BCG strain, we demonstrated that the TSS phenotype is TLR2/TLR4 dependent (5). Therefore, to assess whether the LPS induced TSD phenotype is TLR2/4 dependent, we performed qPCR to evaluate TLR2 and TLR 4 expression. Indeed, we found that LPS treatment led to 2 fold increased expression of TLR2 and TLR4 in EpCAM+/ABCG2+CSCs. (Figure 3D). These results suggest that bacterial product LPS may reprogram EpCAM+/ABCG2+ CSCs to TSD phenotype. Since the TSD phenotype is transient i.e. exhibit rapid expansion for only 1-2 weeks’ time, the LPS-treated ABCG2+ cells were grown for 2-3 weeks and the trypan blue assay was performed to count the viable cells. As shown in Figure 3E, the LPS treated ABCG2+ CSCs showed rapid expansion for the initial 7-days confirming transient expansion of the TSD phenotype. These data indicate that LPS of high grade/relapsed patients 19-22 may selectively reprogram the EpCAM+/ABCG2+CSCs of SCC-25 cell line to the TSD phenotype.

### Assessment of TSD phenotype in the saliva-treated SCC-25 cell line

Saliva is enriched in oral microbiome, which may directly activate the CSC niche defense mechanism in the dormant EpCAM+/ABCG2+ cells of SCC-25. Thus, we reasoned that treating the cancer cell line with oral saliva of patients will lead to internalization of putative TSD phenotype inducing bacteria to EPCAM+/ABCG2+ cells; internalized bacteria will then reprogram these cells to TSD phenotype. To test this hypothesis the saliva samples from n= 25 subjects OSCC that we used to collect LPS (Table 1) were used. The saliva was centrifuged and processed in DMEM culture media, and added to SCC-25 cells. After 3-days, as the cell culture get overwhelmed with bacterial growth, the extracellular bacteria were cleared using amikacin (26) and metronidazole for 24 hours, and then gentamicin was added for the next one week. The treated cells were then cultured for 1 week, and subjected to qPCR gene expression analysis to evaluate the expression of ABCG2, Nanog, and HIF-2alpha, a few key genes involved in the TSD phenotype (26) (Figure 4A). We observed that despite antibiotic treatment, the SCC-25 cells grown with the saliva samples 1-17 remained heavily contaminated with bacterial growth and therefore the cell cultures were discarded. Whereas, patients 18-22 saliva-treated SCC-25 cells exhibited growth. Interestingly, these saliva showed very low number of bacterial 16sRNA load. Thus, we were able to grow saliva-treated SCC-25 cells of patients #18-22 only, whereas the SCC-25 cells grown with the saliva of other patients were heavily contaminated despite antibiotics treatment. Therefore, RNA of SCC-25 cells treated with saliva of these #18-22 patients were obtained for gene expression study of ABCG2, Nanog and HIF-2-alpha, a few genes associated with TSD phenotype, as well as p21, which is downregulated in TSD phenotype. The saliva of two relapsed patients # 21 & 22 exhibited a significant increase in HIF-2 α, Nanog, and ABCG2 compared to untreated cells (Figure 4B). Importantly, in these patients’ treated samples, the EpCAM+/ABCG2+ cells were expanded by 4-7 fold (Figure 5A). These cells exhibited 2-5 fold increased expression of genes involved in TSD phenotype, induction of HIF-2α protein, 4-fold increased invasion in Boyden chamber assay, and 6-fold increase in the transient expansion/proliferation that lasted for two weeks (Figure 5B-C), and suggesting TSD phenotype. These results suggest that the saliva from relapsed patients has the capability to induce expansion of EpCAM+/ABCG2+ CSCs in the SCC-25 cells, and these expanded CSCs exhibited TSD phenotype.

**Figure 4.**
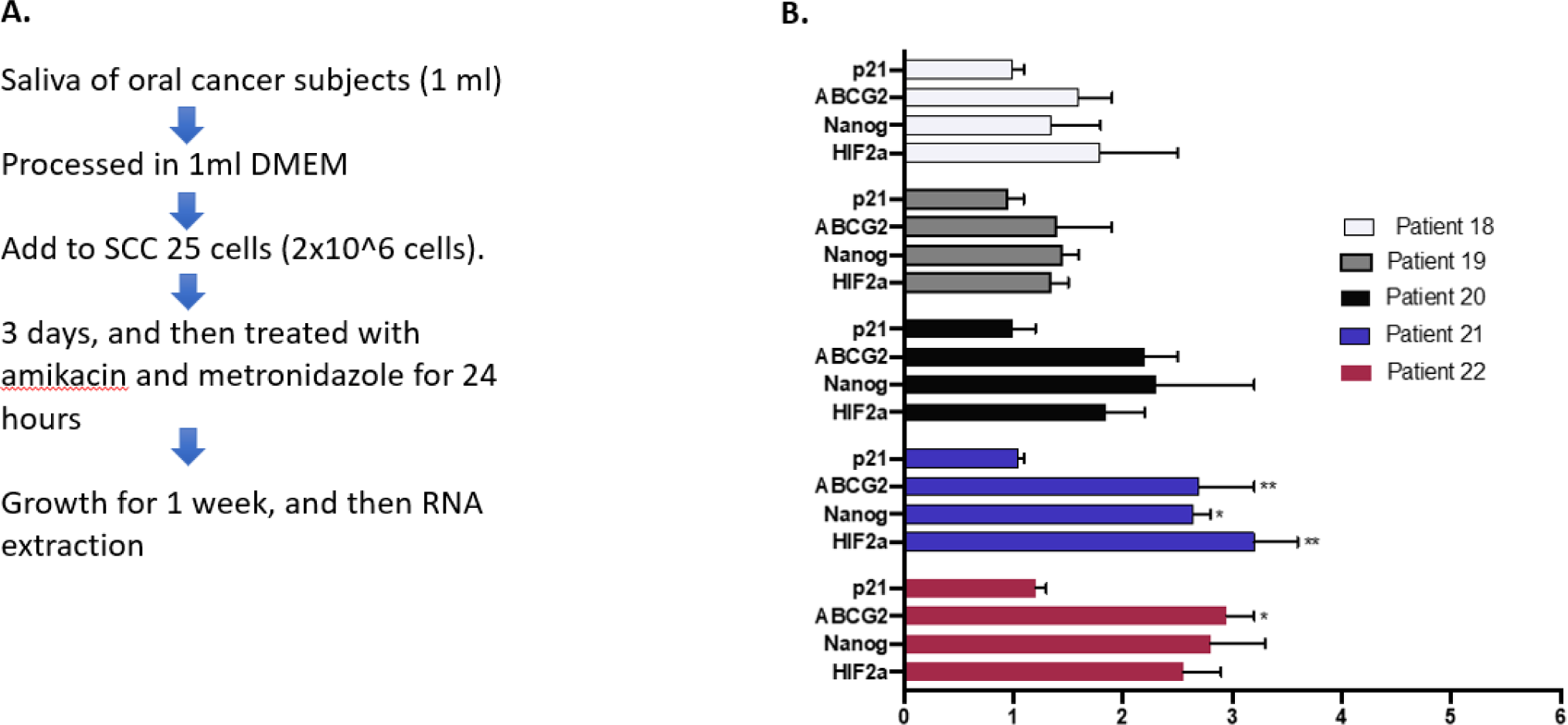
Saliva of oral cancer subjects modulates the genes related to HIF-2α stemness pathway. A) Schematic representation of the experimental design. Saliva from oral cancer subjects is processed in DMEM and added to SCC-25 cells (2×10^6 cells). Three days post treatment, cells in culture were treated with amikacin and metronidazole for 24 hours. Cells were then maintained for 7 days and then assessed for RNA. B) Oral cancer subject’s saliva (table 1) treatment in SCC-25 cells upregulated the genes involved in the HIF-2α stemness pathway. PBS treated cells were used as control in the experiment to compare gene expression. N = 3 independent experiments; error bar represent mean ± SEM. *p<0.05, **p<0.001, ANOVA.

**Figure 5.**
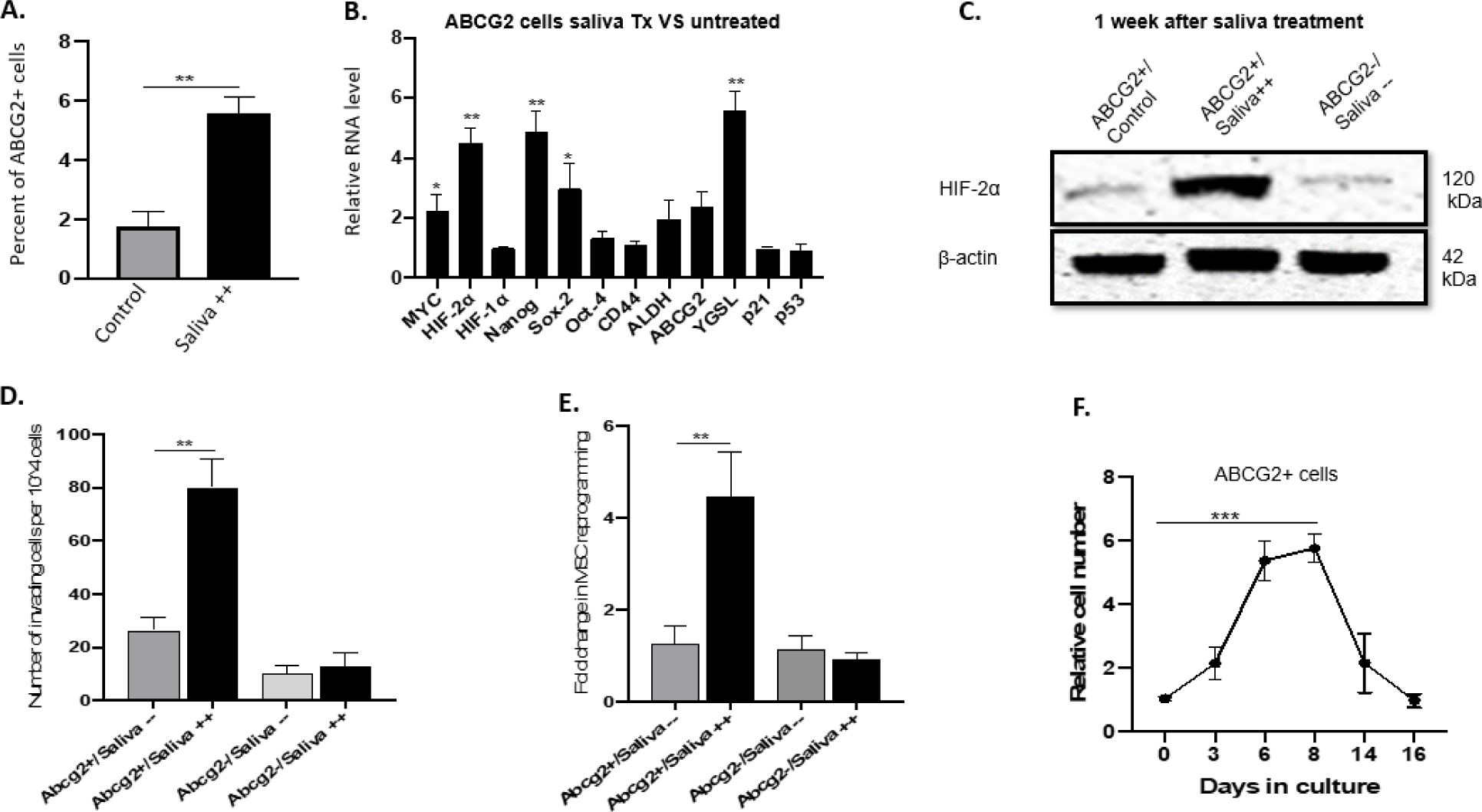
Patient 21 saliva-induced tumor stemness defense phenotypse in SCC-25 cell line. A) Saliva treatment expanded the ABCG2+ population. B) Saliva treatment enhanced the stemness of ABCG2+ cells, with significantly increased expression of genes involved in the HIF-2α stemness pathway. C) Western blot analysis indicates increased protein levels of HIF-2α in ABCG2+ cells of saliva treated group. D) Saliva treatment enhanced invasive potential of ABCG2+ vs ABCG2-cells. E) Saliva treatment enhanced the niche modulatory capacity of ABCG2+ vs ABCG2-cells. F) ABCG2+ cell number increased transiently and lasted for approximately two weeks. N = 3 independent experiments; error bar represent mean ± SEM. *p<0.05, **p<0.001, ***p<0.0001, ANOVA.

To evaluate if the saliva-induced TSD phenotype is TLR4 pathway-dependent, we performed qPCR analysis of TLR2 and TLR4 gene expression. We found that the saliva-treated ABCG2+ cells exhibited a significant increase in TLR4 expression as compared to the controls (Figure 6A). We then evaluated the number and percentage of EpCAM+/ABCG2+ cells with and without the addition of anti-TLR2/4 antibodies (Invivogen) (neutralizing antibody against TLR2/4). Notably we found that the addition of anti-TLR4 resulted in decreased number and percentage of ABCG2+ cells, while the addition of anti-TLR2 had no significant effect on that cell population (Figure 6 B&C). We also performed qPCR analysis of HIF-2α and p21 gene expression with and without the addition of anti-TLR4. We have found that the addition of anti TLR4 resulted in no increase in the expression of HIF-2α, or the decrease in p21 (Figure 6D). These results suggest that the saliva induced stemness is TLR4 pathway dependent.

**Figure 6.**
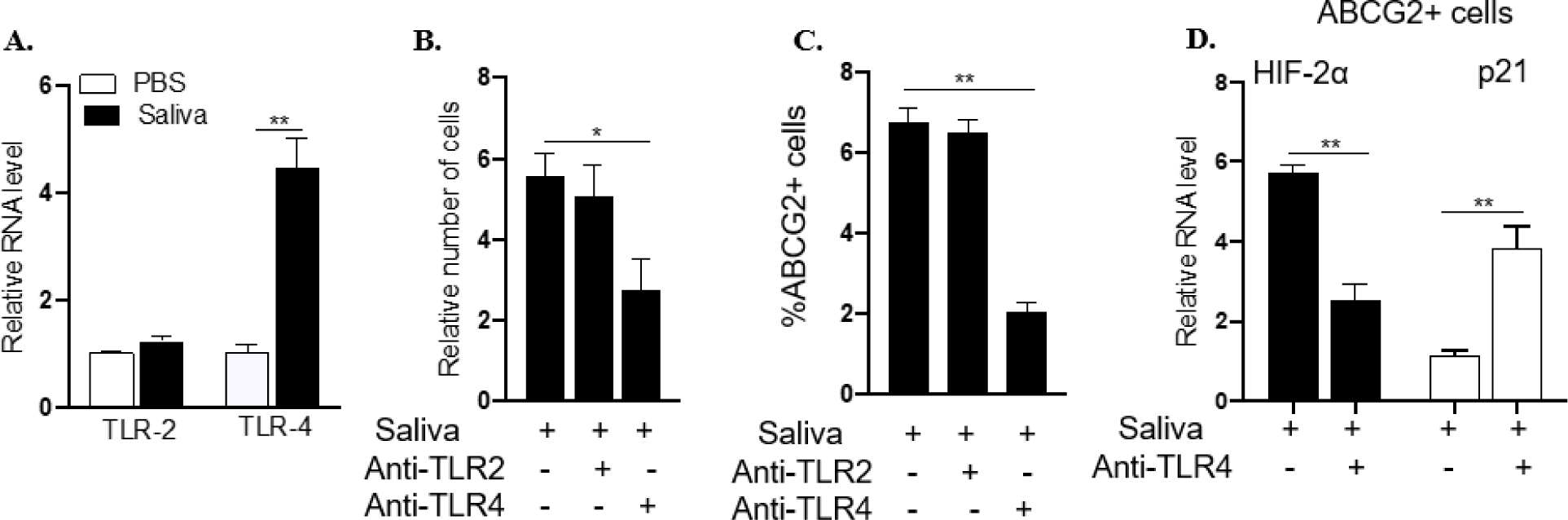
Patient 21 saliva induced tumor stemness is TLR4 pathway-dependent. A) Day 7 after saliva treatment. qPCR analysis indicates that the saliva treatment increased TLR4 expression. B) Relative number of ABCG2+cells following treatment with anti-TLR2/4 antibodies (Invivogen) (neutralizing antibody against TLR2/4). C) Percentage of ABCG2+ cells following treatment with anti-TLR2/4 antibodies D) qPCR analysis of HIF-2α and p21 gene expression in saliva treated group with and without anti TLR4 indicating that the saliva induced stemness is TLR4 pathway dependent. N = 3 independent experiments; error bar represent mean ± SEM. *p<0.05, **p<0.001, ANOVA.

### Live bacteria are needed for the saliva induced TSS phenotype

To investigate whether the live bacteria in the saliva are needed for the induction of TSD phenotype in SCC-25 cells, the processed saliva samples of patients 21 &22 were treated with broad-spectrum antibiotics ciprofloxacin (50 ug/ml dose for 3 days), and metronidazole (10ug/ml) to kill the live bacteria. The culture treated saliva was then added to the growing SCC-25 cells; these cells were then subjected to TSD phenotype analysis. As shown in Figure 7, the ciprofloxacin/metronidazole treated saliva exhibited no increase in the expression of MYC, HIF-2 α, Sox-2 or Nanog suggesting that live bacteria is needed for TSS phenotype induction.

**Figure 7.**
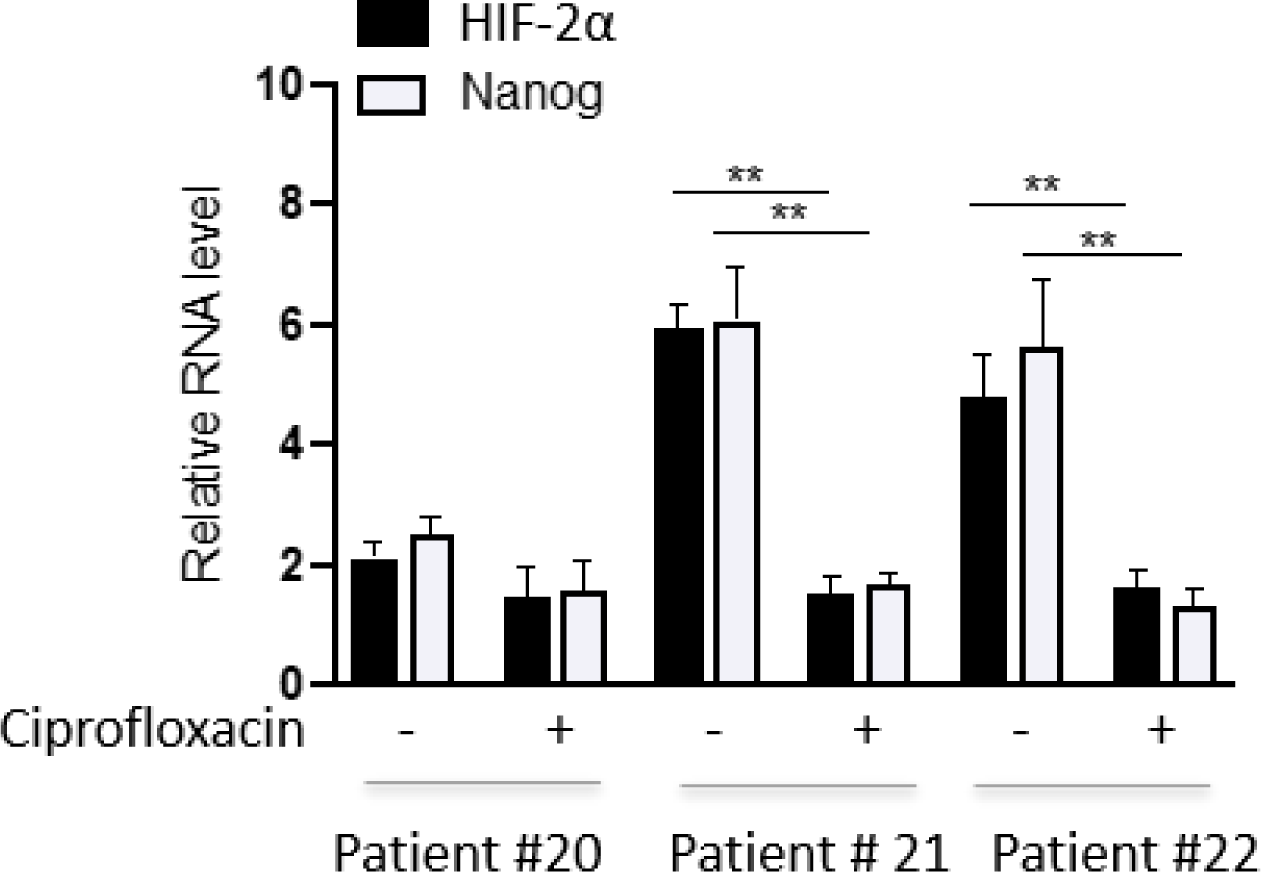
Ciprofloxacin treated saliva from patient 20, 21 and 22 effect on SCC-25 cells. qPCR analysis of ABCG2+ cells immunomagnetically sorted from SCC-25 cells pre-treated with saliva from Patient 20, 21 and 22, with and without ciprofloxacin for 7 days and evaluated for expression of HIF-2α and Nanog. Data was normalized to GAPDH to obtain the ΔΔCT value as previously described. N = 3 independent experiments; error bar represent mean ± SEM. **p<0.001, ANOVA.

### ABCG2+ cells exhibit the presence of *F.nucleatum*

To investigate whether specific bacteria may be involved in the reprogramming of SCC-25 cells, we obtained the cell extracts from the ABCG2+ cells with TSD phenotype, and also from the ABCG2-cells; the lysed extract was then cultured in both aerobic and anerobic bacterial culture. We observed growth in the anerobic culture having a morphology of F. nucleatum. We confirmed the phenotype by performing RT-PCR with an *F.nucleatum* specific primer (Figure 7A-B). The results suggests that F. nucleatum internalizes within the SCC-25 cells and may modulate the EpCAM+/ABCG2+ cells to TSD phenotype. Indeed, the co-culture of SCC-25 cells with *F.nucleatum* isolated from the saliva treated EpCAM+/ABCG2+ cells (Figure 8A) led to the expansion of dormant EpCAM+/ABCG2+ cell fraction of SCC-25 (Figure 8B). These cells showed increased clonogenicity, and increased levels of TSD phenotype associated MYC, HIF-2α and Nanog genes and corresponding protein levels (Figure 8 C-D). Additionally, the addition of anti-TLR4 results in downregulation of HIF-2α expression, and upregulation of p21 expression (Figure 8E) suggesting that *F nucleatum* induced stemness is TLR 4 dependent.

**Figure 8.**
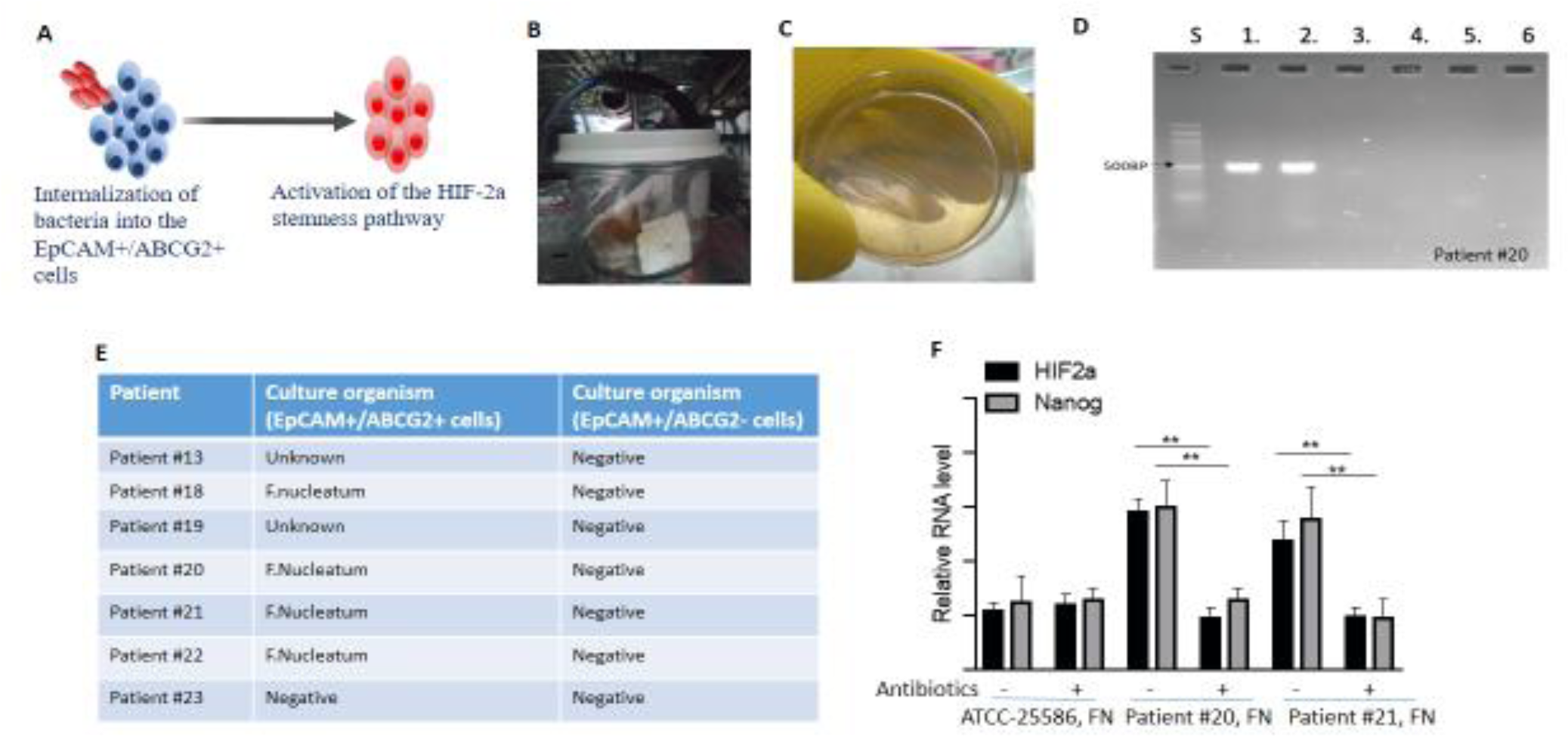
SCC-25 cells derived ABCG2+ cells treated with Patient 4 saliva is f-nucleatum positive. A) The figure depicts internalization of bacteria into the EpCAM+/ABCG2+cells and consequent activation of HIF2α stemness pathway B) Selective culture of FN in anaerobic jar C) Growth of colonies in Fusobacterium egg agar plates.D) Agarose gel electrophoresis of the amplification products obtained by the F.Nucleatum was performed. Each (4 ng) of the bacterial genomic DNA was used as the PCR template. Lanes: S, 50 base pair DNA ladder (Hi Media.); 1. F. nucleatum ATCC 25586; 2. ABCG2+ cells of Patient #20 saliva. 3. Control SCC-25 cells 4. ABCG2-cells of Patient #20 saliva. 5. Sterilized deionized water (negative control). The filled arrowhead indicates the PCR products of F. nucleatum (495 bp) E) List of patients from whom presence FN was found in ABCG2+ cells F) Relative RNA level of HIF 2α and nanog in culture of FN from ATCC 25586,Patient 20 and Patient 21in presence of antibiotics and without antibiotics.

**Figure 9.**
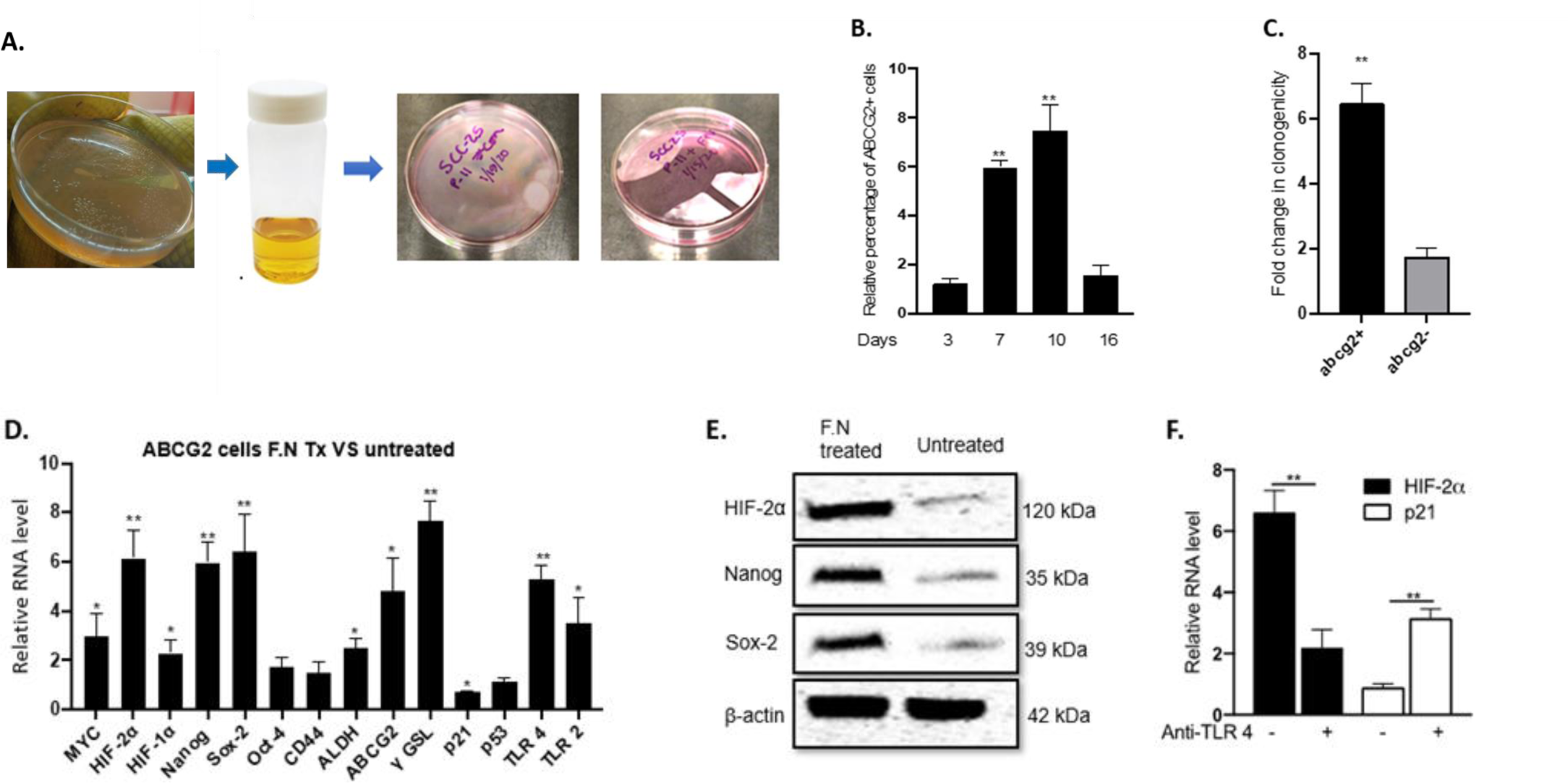
F-nucleatum induces tumor stemness switch in ABCG2+ cells. A) F.Nucleatum (ATCC:25586) were cultured in Fusobacterium egg agar plates for 1 week and then sub cultured in brain-heart infusion broth for 24 hours were added to SCC-25 culture in MOI 20. After 3 days, cells were treated with antibiotics amikacin and Metronidazole for 24 hours, and then incubated for two weeks.,B) Relative percentage of ABCG2+ cell after treatment with FN C) F. nucleatum infection led to expansion of ABCG2+ cell population vs ABCG-cells. D) qPCR analysis indicates increased expression of genes involved in the HIF-2α stemness pathway. E) Western blot analysis indicates increased protein levels of HIF-2α, Nanog and Sox-2 in the F. nucleatum treated group. F) qPCR analysis of HIF-2α and p21 gene expression in F. nucleatum treated group with and without anti TLR4 (neutralizing antibody against TLR4) indicating that the F. nucleatum induced stemness is TLR4 pathway dependent. N = 3 independent experiments; error bar represent mean ± SEM. *p<0.05, **p<0.001, ANOVA.

### Primary EpCAM+/ABCG2+ CSCs of relapsed patient contain intracellular *F.nucleatum*

To investigate if primary tumor-derived EpCAM+/ABCG2+ cells contain intracellular bacteria, we obtained primary tumors from n=10 OSCCs including the primary tumors of relapsed subjects (n=5). The immunomagnetically sorted EpCAM+/ABCG2+ cells were lysed, and subjected to in vitro anaerobic culture. Then, the presence of F.N was confirmed by qPCR. The rest of the EpCAM+/ABCG2-cells were also used to obtain the *F.nucleatum* culture. We were able to isolate *F.nucleatum* from n=3 relapsed tumors. The isolated *F.nucleatum* demonstrated the induction of TSD phenotype in the SCC-25 cells.

### CSC-derived FN exhibit phenotypic changes

Two outer membrane proteins found in *F.nucleatum*, specifically fibroblast activation protein 2 (Fap2) and Fusobacterium adhesin A (FadA), have recently garnered attention. Fap2 is a protein with a sensitivity to galactose, acting as both a hemagglutinin and an adhesive protein. It plays a crucial role in *F.nucleatum’s* ability to invade human cells by binding to D-galactose-β (1–3)-N-acetyl-D-galactosamine (Gal-GalNAc)(27) (28). FadA is a surface adhesive protein found exclusively in *F. nucleatum*, and it plays a critical role in facilitating the process of cell-to-cell attachment. (29). These two mentioned proteins have significantly contributed to host cell attachment, which in turn influences the modulation of signaling pathways associated with colorectal cancer (CRC) (29) (30). Therefore, we investigated whether the F.N bacteria may undergo phenotype changes, specially the increase of FADA and FAP2 activities as investigated by congo red staining method. The FN bacteria recovered from the primary tumors and also from the saliva treated SCC-25 ABCG2+ cells were lysed and subjected to congo red assay, as well PCR for FaDA gene. The results show that the FN of EpCAM+/ABCG2+ obtained either from primary tumors or the F.N. infected SCC-25 cells but not patient-derived saliva express high FaDA gene, and exhibit marked congo red absorption. We also found that ATCC derived *F.nucleatum* grown in anaerobic media did not exhibit marked congo red absorption. However, intracellular ATCC-derived *F.nucleatum* recovered from the EpCAM+/ABCG2+ cells of SCC-25 showed the congo red staining and the expression of FADA and FAP2 gene. These results indicate that a host-pathogen interaction between CSCs and *F.nucleatum* may occur that confer phenotypic changes to *F.nucleatum*.

## Discussion

Oral microbiota may induce stemness by activating inflammatory pathways. Whether the microbiota may also induce CSC niche defense is not studied. Here, we provided the first experimental approach to studying the microbiota-induced CSC niche defense. We found that LPS treatment obtained from the saliva of OSCC subjects induces a generalized upregulation of genes involved in tumor stemness. However, only in 4/25 OSCC subjects 10 ml of saliva contain enough LPS to induce stemness, indicating that LPS is the most unlikely major player of microbiota induced stemness. Whereas, four subjects with relapsed OSCC cases showed the presence of very low LPS in the saliva, but were able to induce TSD phenotype. FN present in the saliva of these patients internalized into the ABCG2+ cells leading to the activation of CSC niche defense. In these subjects, we recovered live FN from the EPCAM+/ABCG2+ cells of SCC-25. Notably, 3/5 OSCC relapsed tumor derived EpCAM+/ABCG2+ contained *F.nucleatum*.

Han *et al* for the first time demonstrated the internalization of FN inside the human gingival epithelial cells (HGEC) which led to increased production of IL-8 a proinflammatory cytokine (31). Notably another study reported increased production of IL8, MMP 1 and MMP 9 in OQ01 (primary head and neck cancer cell line) infected by live FN bacteria. The increased secretion of IL 8 led to cancer cell invasion (7). Intracellular *F.nucleatum* may induce inflammatory pathways leading to the induction of IL-8 secretion.

These inflammatory pathways may enhance the MYC-HIF-2alpha stemness pathway for reprogramming of CSC to TSD phenotype(14). In this context, we reasoned that live bacteria *Fusobacterium nucleatum* in saliva is involved in inducing CSC niche defense via inflammatory pathway. Notably we used ciprofloxacin to treat saliva and then treat SCC 25 cells and we found that ciprofloxacin treatment led to low expression of HIF 2α, and Nanog suggesting that live bacteria is needed for the induction of TSD phenotype. Moreover, direct FN infection of SCC 25 cells led to increased expansion of EpCAM+/ABCG2+ CSCs exhibiting TSD phenotype. Thus, FN may induce a host-pathogen interaction, where the host cells may activate inflammatory pathways to defend their niches against the invading bacteria. Notably, we have reported that CSC defend their niche via TLR2/TLR 4 pathway following BCG infection (5). In this context, CSC niche defense against FN is also TLR 4 dependent. When anti TLR 4 was used to treat the EpCAM+/ABCG2+ CSC infected with F.N., the expression of HIF 2α was markedly reduced, while p21 expression was upregulated indicating the differentiation of the TSD phenotype. Future research is needed to investigate how a few intracellular F.N. may activate inflammatory and stemness pathways to reprogram CSCs to TSD phenotype.

Previously we have reported that chemotherapy, oxidative stress prevailing in the CSC niche can reprogram the CSCs to TSD phenotype (19) (5, 32). Moreover we have also reported that internalization of Mycobacteria in CSCs of TSD phenotype can induce PIBA in apoptosis in other CSCs in the niche (5). However our results suggest that internalization of FN in the ABCG2+ CSCs in the SCC 25 cells activates the CSC niche defense with high expressions of embryonic stemness genes, high clonogenicity of ABCG2+ and high invasive ability. Thus the results infer that FN has pro tumorigenic ability rendering OSCC to be aggressive. Importantly the results presents us with new avenue to study bacteria induced aggressiveness in oral cancer and bacteria can be biomarker in OSCC.

Our findings on LPS mediated TSD phenotype need critical review. One of the key biomarker in OSCC can be detection of bacterial endotoxin such as LPS in the saliva of oral cancer patients. LPS biosynthesis has been reported in OSCC sites and also has been reported to induce OSCC progression and migration (6). Oral bacteria such as *P.multocida* are dependent on LPS for infection (33). Importantly LPS induces inflammatory cytokines which in turn plays key role in cancer cell invasion, tumor progression and metastasis. A study has reported secretion LPS via TLR 4 may induce stemness in oral cancer cells by activating inflammatory pathways(9). In this context our results suggest that LPS induces TSD phenotype in SCC 25 cells via inflammatory pathway. LPS treatment led to expression of stemness genes with marked expression of ABCG2+ CSC marker. Also, LPS treated ABCG2+ CSCs showed high clonogenicity. Notably invasion assay showed that LPS treated ABCG2+ CSCs had high expression of IL 10 and these CSCs showed the expression of TLR2/TLR4. Thus LPS has the ability to induce TSD phenotype and can be a biomarker of aggressive TSD phenotype in OSCC. However, salivary LPS can also be released from a number of other oral bacteria in the oral cavity of OSCC patients. In this context LPS from specific pathogenic oral bacteria associated OSCC can play a specific role in inducing TSD rather than LPS from other oral bacteria. Therefore this study has the limitation of how LPS from specific pathogenic oral bacteria can induce TSD phenotype in OSCC patients. Notably, we reason that intracellular bacteria can be specific biomarker in OSCC patients.

Studies have shown that bacteria resides in tumor tissues. Indeed Hooper *et al* found heterogeneous viable bacterial population in the cancer cells and TME of primary OSCC using 16s RNA sequencing (34). Notably another study showed the presence of bacteria and LPS intracellular to seven solid tumors types. However no study has shown bacteria intracellular to CSCs. In our study we have shown for the first time how internalization of FN in ABCG2+ CSCs of OSCC reprogram the tumors to highly tumorigenic TSD phenotype. In this context, the presence of intracellular bacteria in primary OSCC can be a potential biomarker of aggressive OSCC.

### Conclusion

Oral squamous cell carcinoma is highly aggressive cancer. The host pathogen interaction of oral bacteria FN and OSCC can reprogram the cancer to a highly aggressive phenotype. Importantly, earlier studies have shown the role of oral bacteria in OSCC invasion and progression but these studies have not shown the innate defense mechanism and internalization of bacteria into the CSCs of OSCC. Here we have shown how internalization of FN into the CSCs activate defense mechanism of enhanced stemness of TSD phenotype. In the *in-vitro* we have efficiently shown that LPS from oral bacteria can regulate stemness via TLR 4 dependent pathway. Notably we were able to show that patient saliva has pathogenic oral bacteria FN which can internalize in the CSCs of OSCC and reprogram OSCC to a highly tumorigenic phenotype. In conclusion it is clear that our *in-vitro* model of host pathogen interaction is potential model to study oral bacteria stem cell niche defense mechanism and also can be potential model to study the oral bacteria as a biomarker of OSCC aggressiveness.

## Data availability statement

The raw data supporting the conclusions of this article will be made available by the authors upon reasonable request.

### Ethics statement

This study was reviewed and approved by the Institutional Ethics Committee of KaviKrishna Laboratory, and Thoreau Laboratory for Global Health, M2D2, University of Massachusetts, Lowell, MA, USA. Necessary patient consent was taken as per guidelines of the Institutional Ethics Committee of KaviKrishna Laboratory, IIT Guwahati Research Park, Indian Institute of Technology, Guwahati, India.

## Author Contributions

BD has designed the experiment model, analyzed the data, and edited the article. PJS has done the sample collection, and experiments, written, edited, and proofread the article and also created the figures and table. LP has done experiments, and written and edited the article. SM has done experiments, and written and proofread the article. TB and RD have collected saliva samples and contributed to the patient’s care. BP has performed the experiments and analyzed the data. IA has performed the experiments All authors contributed to the article and approved the submitted version.

## Acknowledgements

We sincerely extend our gratitude to the members of KaviKrishna laboratory, IIT Guwahati Research Park, Indian Institute of Technology, Guwahati, India and Thoreau Laboratory for Global Health, M2D2, University of Massachusetts, Lowell, MA, USA who provided insight and expertise that greatly helped in the completion of this research article.

